# Performance of an IAVI-African Network of Clinical Research Laboratories in Standardized ELISpot and Peripheral Blood Mononuclear Cell Processing in Support of HIV Vaccine Clinical Trials

**DOI:** 10.1101/300087

**Authors:** Robert Langat, Bashir Farah, Jackton Indangasi, Simon Ogola, Gloria Omosa-Manyonyi, Omu Anzala, Jean Bizimana, Emmanuel Tekirya, Caroline Ngetsa, Moses Silwamba, Enoch Muyanja, Paramesh Chetty, Maureen Jangano, Nancy Hills, Jill Gilmour, Len Dally, Josephine H. Cox, Peter Hayes

**Affiliations:** Kenya AIDS Vaccine Initiative -Institute of Clinical Research, University of Nairobi, Nairobi, Kenya; International AIDS Vaccine Initiative (IAVI) Human Immunology Laboratory, Imperial College, London, United Kingdom; Projet San Francisco, Kigali, Rwanda; Kenya Medical Research Institute Centre for Geographical Medicine Research Coast, Kilifi, Kenya; Zambia EMORY HIV Research Project, Lusaka, Zambia; Ugandan Virus Research Institute-IAVI, Entebbe, Uganda; IAVI,Johannesburg, South Africa; Clinical Laboratory Services, Johannesburg, South Africa; University of California, San Francisco; International AIDS Vaccine Initiative, New York, New York, United States of America; Emmes Corporation, Rockville Maryland, USA; Clinical Trials Program, Vaccine Research Center, National Institutes of Health, Bethesda, MD, USA

## Abstract

Immunological assays performed in different laboratories participating in multi-centre clinical trials must be standardized in order to generate comparable and reliable data. This entails standardized procedures for sample collection, processing, freezing and storage. The International AIDS Vaccine Initiative (IAVI) partnered with local institutions to establish Good Clinical Laboratory Practice (GCLP)-accredited laboratories to support clinical trials in Africa, Europe and Asia. Here we report on the performance of seven laboratories based in Africa and Europe in the interferon-gamma enzyme-linked immunospot (IFN-γ ELISpot) assay and peripheral blood mononuclear cell (PBMC) processing over four years. Characterized frozen PBMC samples from 48 volunteer blood packs processed at a central laboratory were sent to participating laboratories. For each stimulus, there were 1751 assays performed over four years. 98% of these ELISpot data were within acceptable ranges with low responses to mock stimuli. There were no significant differences in ELISpot responses at five laboratories actively conducting immunological analyses in support of IAVI sponsored clinical trials or HIV research. In a separate study, 1,297 PBMC samples isolated from healthy HIV-1 negative participants in clinical trials of two prophylactic HIV vaccine candidates were analysed for PBMC yield from fresh blood and cell recovery and viability following freezing and thawing. 94 % and 96 % of samples had fresh PBMC viabilities and cell yields within the pre-defined acceptance criteria while for frozen PBMC, 99 % and 96 % of samples had acceptable viabilities and cell recoveries respectively, along with acceptable ELISpot responses in 95%. These findings demonstrate the competency of laboratories across different continents to generate comparable and reliable data in support of clinical trials.

**Importance:** There is a need for the establishment of an African network of laboratories to support large clinical trials across the continent to support and further the development of vaccine candidates against emerging infectious diseases such as Ebola, Zika and dengue viruses and the continued HIV-1 pandemic. This is particularly true in sub-Saharan Africa where the HIV-1 pandemic is most severe. In this report we have demonstrated by using standardized SOPs, training, equipment and reagents that GCLP-accredited clinical trial laboratories based in Africa and Europe can process clinical trial samples and maintain cell integrity and functionality demonstrated by IFN-γ ELISpot testing, producing comparable and reliable data.

## Introduction

Clinical trials related to HIV, malaria and tuberculosis (TB) have been conducted in Africa for decades (1–4). In order to harmonize the immunological data generated from these clinical trials, laboratories responsible for clinical sample processing must establish standardized procedures to meet International Conference on Harmonization (ICH) Good Clinical Practice (GCP) and World Health Organisation (WHO) guidelines for collection, processing and storage of samples and for immunological assays. The International AIDS Vaccine Initiative (IAVI) has partnered with local institutions and established Good Clinical Laboratory Practice (GCLP)-compliant laboratories across Africa, Europe and India to conduct safety and immunogenicity assessments in support of clinical trials of HIV vaccine candidates (5, 6). These laboratories are equipped to process and store samples for later testing and are able to perform ELISpot and flow cytometry immunological assays. IAVI has conducted over 20 phase 1 HIV vaccine trials (www.iavi.org/trials-database), with the majority in Africa (7–10). To ensure uniformity of data, IAVI sponsored a central laboratory at Imperial College London (Human Immunology Laboratory) to provide Standardised Operating Procedures (SOPs), training, critical assay reagents, long-term centralised sample storage and to perform immunological assays where local laboratories have no capability. IAVI and partners have developed the capacity of local personnel professionally and academically, through technical training, mentoring and funding for investigator-initiated research projects. These GCLP-accredited-laboratories have also been used as reference laboratories allowing local (and International) research organizations to utilize existing facilities for their research and staff training in technical assays and GCLP guidelines.

The performance of the IFN-γ ELISpot assay across multiple laboratories both within and across continents is critical to the generation of standardized data on vaccine immunogenicity (11). Previous studies (12) showed varied responses across laboratories while more recent ELISpot proficiency studies (13–15) have shown significantly improved results. These recent findings demonstrated that GCLP-accredited laboratories participating in proficiency testing over time can generate highly concordant results.

The majority of laboratories performing end-point ELISpot assays and enrolled in ELISpot proficiency schemes are based in Europe and USA (12, 15, 16). With increased focus on testing HIV vaccine products where the pandemic is most severe and with renewed interest in many “orphan” tropical infectious diseases (17), development of laboratory networks able to support large clinical trials across sub-Saharan Africa has gained importance. In order for IAVI-supported laboratories to meet international standards, they were enrolled into an IAVI-sponsored ELISpot proficiency scheme organised by Clinical Laboratory Services, South Africa.

## Methods and Materials

### Participating laboratories

All participants were IAVI-supported laboratories including: 1) Kenya AIDS Vaccine Initiative-Institute of Clinical Research (KAVI-ICR) University of Nairobi, Nairobi, Kenya; 2) IAVI Human Immunology Laboratory (HIL), Imperial College London, United Kingdom; 3) Uganda Virus Research Institute (UVRI)-IAVI, Entebbe, Uganda; 4) Clinical Laboratory Services (CLS), Witwatersrand University, Johannesburg, South Africa; 5) Kenya Medical Research Institute Centre for Geographical Medicine Research Coast (KEMRI-CGMRC), Kilifi, Kenya; 6) Zambia EMORY HIV Research Project (ZEHRP), Lusaka, Zambia; and 7) Projet San Francisco (PSF), Kigali, Rwanda. The laboratories were located in six countries (Kenya, Uganda, Rwanda, Zambia, South Africa and UK). CLS is not supported by IAVI but provide PBMC, pretesting and shipping services for the ELISpot proficiency testing. CLS participate in quarterly testing of the ELISpot proficiency and are included in this analysis.

## Laboratory Preparation

### Process of establishing clinical trial laboratories under GCLP guidelines

Comprehensive training programs, standardized GCLP-compliant serviced and calibrated equipment and quality control (QC) measures were integral to establishing IAVI’s laboratories. The key elements for establishing the clinical trial laboratories are shown in **Table 1** and described in detail in the Supplementary Information.

**Table 1:**
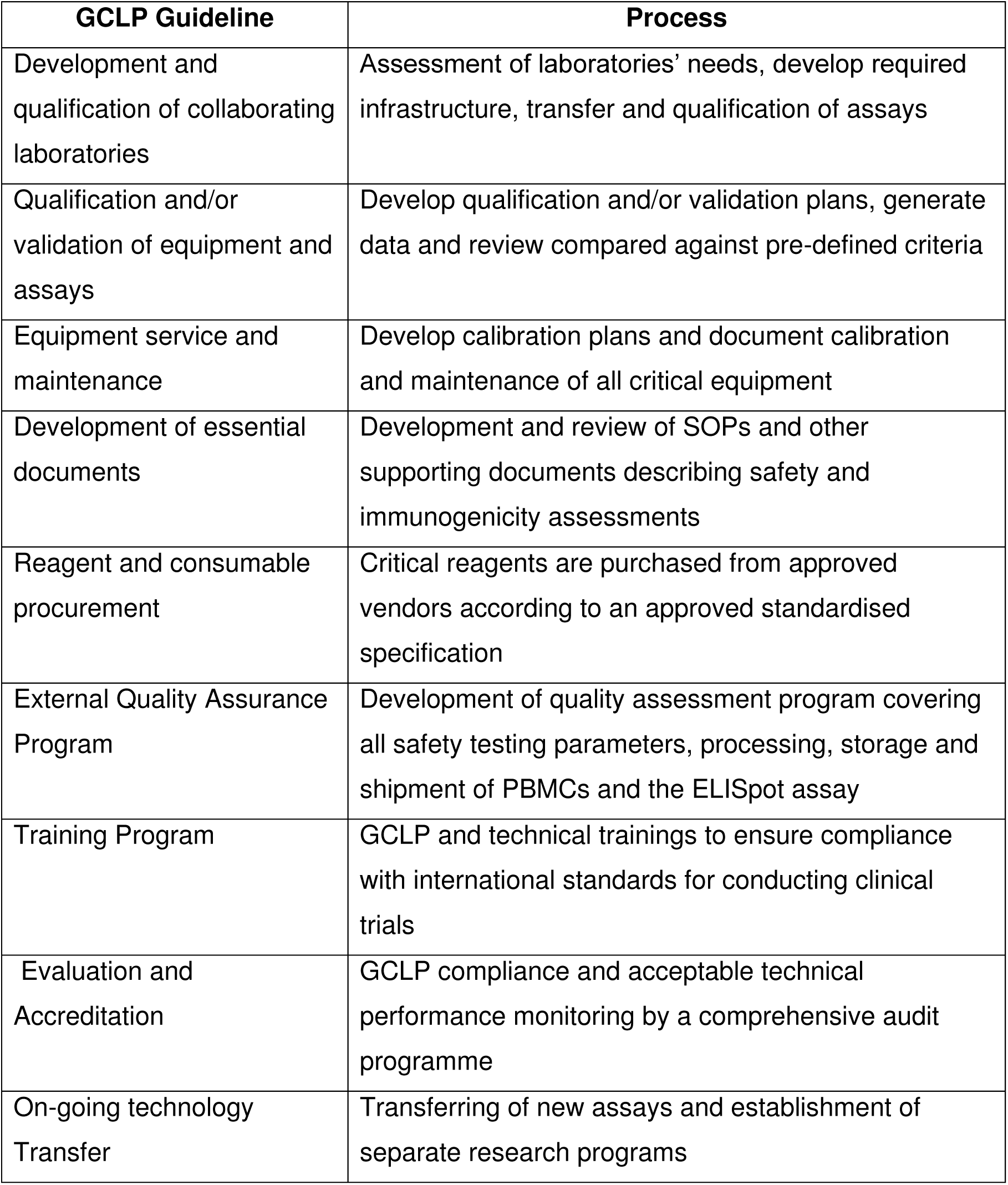
Summary of the process of establishing clinical trial laboratories under GCLP guidelines.

The quality systems and SOPs were designed to minimise failure, identify problems, initiate corrective actions and monitor resolutions. Two laboratories (HIL and CLS) have been designated as support and QC management. London (United Kingdom) and Johannesburg (South Africa) were ideal locations to support a global clinical trial program. Both are major international hubs in Europe and Africa, with direct flights to and from the above IAVI-supported laboratories, thereby reducing time, cost and risk to samples in transit.

### ELISpot proficiency panel design

IAVI–GCLP laboratories were enrolled in an ELISpot proficiency scheme coordinated by CLS, South Africa. At CLS, PBMC samples from 48 volunteer blood packs were processed and shipped to IAVI-supported laboratories for assessment of cell viability and IFN-γ ELISpot responses. Two peptide pools were used; a pool of 32 8–10mer peptides representing immunodominant CD8+ T-cell epitopes from Cytomegalovirus, Epstein Barr virus and Influenza virus (CEF) (18) and a pool of 138 15mer peptides overlapping by 11 amino acids spanning the human Cytomegalovirus (CMV) pp65 protein at a final assay concentration of 1.5 μg/mL per peptide. A positive control of phytohaemagglutinin (PHA-L; Sigma L4144) at 10μg/mL and a mock stimulus (medium / dimethyl sulfoxide (DMSO) alone) were used.

### PBMCs for ELISpot Proficiency

Each ELISpot proficiency panel consisted of 6 PBMC samples sufficient for monthly testing of the same 6 samples for 6 months at all laboratories. Different PBMC batches were provided from CLS each 6-month period over four years as shown in Figure 1. Four laboratories; KAVI-ICR, UVRI-IAVI, PSF-Kigali and the HIL actively performing immunological assays in support of IAVI clinical trials, conducted monthly ELISpot testing while 3 laboratories; KEMRI-CGMRC, ZEHRP and CLS not actively performing immunological assays conducted quarterly ELISpot testing. Raw ELISpot data were submitted to the IAVI HIL central data repository for evaluation of responses and results posted onto an access-restricted CLS website.

**Figure 1.**
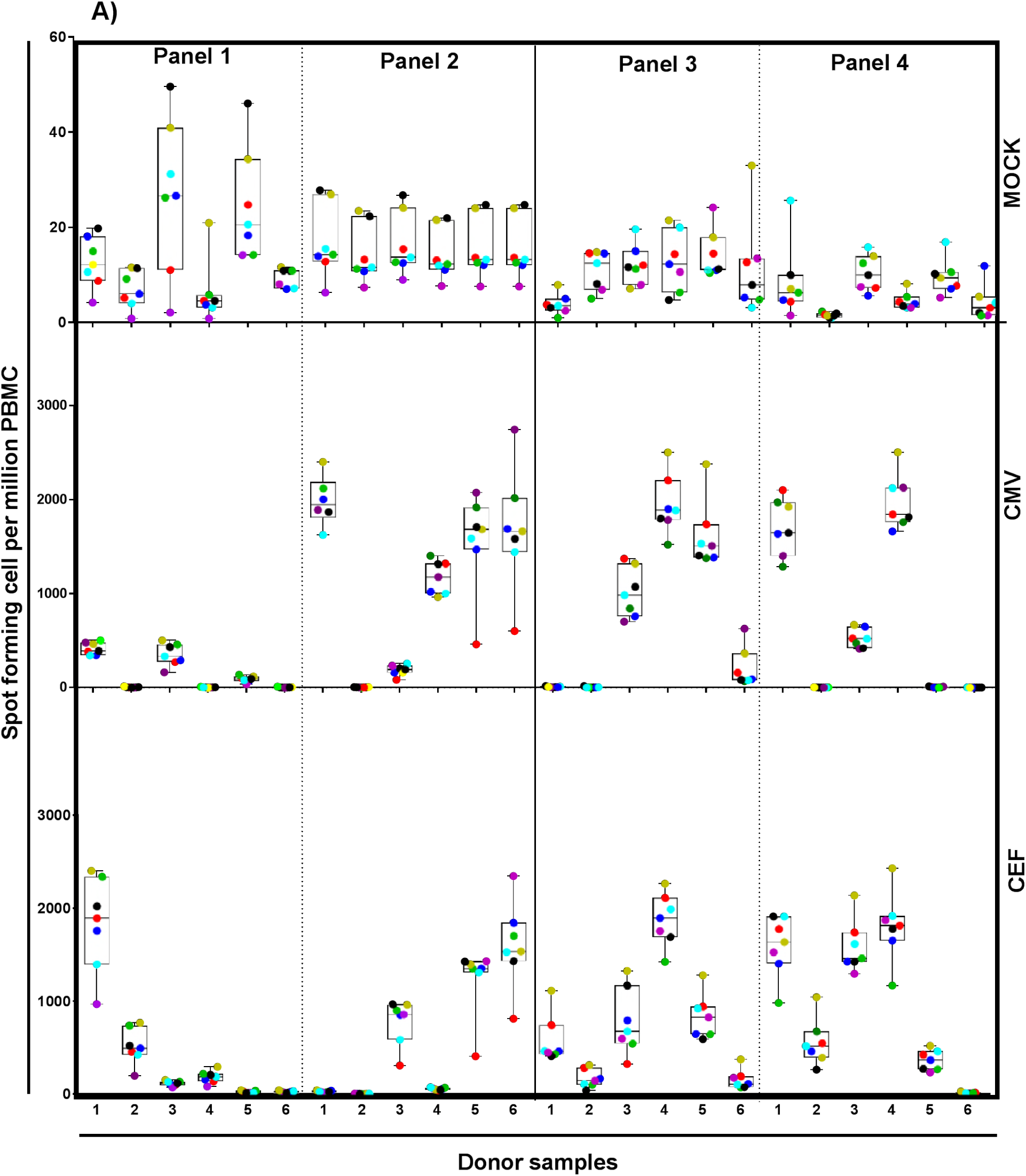
ELISpot spot forming cells (SFC) per million PBMC and variability across laboratories represented in *box plots*. **A)** years 1-2 and **B)** years 3-4. Each panel consists of the same 6 donor samples tested over a 6-month period by 7 laboratories. Box plots represent the quartiles, horizontal line the median and whiskers represent the maximum and minimum values. Each point represents average SFC/10^6^ PBMC from replicates per donor at each laboratory. Laboratories are color-coded as follows: Red = KAVI-ICR, Blue = UVRI-IAVI, Green = PSF, Purple = ZEHRP, Yellow =KEMRI-CGMRC, Cyan =CLS and Black = HIL.

### Sample preparation and immunological assessment

Minimal Information About T-cell Assays (MIATA) guidelines were followed for transparent reporting on immunological assays (19).

#### PBMCs and clinical trial samples

PBMCs used in ELISpot proficiency testing were prepared by Ficoll density centrifugation of buffy coats obtained from 48 healthy, HIV-1 negative blood donors (South African National Blood Bank, Johannesburg, South Africa). Clinical trial samples were obtained from heparinized blood from healthy HIV-1 negative placebo and vaccine recipients participating in clinical trials of two prophylactic HIV vaccine candidates at KAVI-ICR, UVRI-IAVI, PSF and ZEHRP (8, 10). Isolated PBMCs had to meet the following pre-defined acceptance criteria: 1) viability above 90% and cell yield greater than 0.7 ×10^6^ PBMC/mL blood for freshly isolated PBMCs, 2) viability above 80% for frozen PBMCs following thaw and overnight rest and 3) processing of PBMCs from blood draw to start of freezing to be completed within 6 hours. Cell counts and viabilities were determined using a Vi-Cell XR (Beckman Coulter, UK) automated cell counter. PBMCs were frozen in freezing media containing 10% DMSO diluted in fetal calf serum (FCS) using a rate-controlled freezer (Planer, Sunbury-On-Thames, UK) which lowered the temperature by 1°C per minute from +4°C to -80°C then 10°C per minute to -120°C, before transfer to vapour phase liquid nitrogen (LN) until assayed. PBMC for ELISpot proficiency were stored in vapour-phase liquid nitrogen prior to shipment to participating laboratories using temperature-monitored cryogenic dry shippers (MVE Jencons, United Kingdom). The shippers were calibrated according to an SOP. Briefly, empty dry shippers were pre-weighed before filling with LN then left overnight for adsorption of LN and decanted the following day. Shipper weight loss and temperatures were recorded over the next 5 days. The dry shippers were also fitted with the temperature loggers where temperature data were downloaded upon receipt of the dry shippers at the participating laboratories. In order for the dry shippers to pass the calibration, the average 24-hour weight loss over the 5-day calibration was 0.6kg+/-10% and temperature <-190°C. Following receipt, PBMC were transferred to vapour-phase liquid nitrogen until assayed.

#### IFN-γ ELISpot Assay

Cell recovery and viability of samples thawed and rested overnight for ELISpot testing were analysed. PBMCs were removed from LN storage and transported to the laboratory in dry ice and immediately immersed in a 37°C water bath until a small amount of ice remained. Cells were transferred to 10mL cell culture medium (RPMI 1640 supplemented with 10% heat-inactivated FCS, 1 mM L-glutamine, 100 units/mL penicillin, 100 μg/mL streptomycin, 1mM sodium pyruvate and 0.5 mM HEPES (R10)), centrifuged at 250g/10 mins, supernatants decanted, cell pellets disrupted and resuspended in 4mL RPMI 1640 supplemented with 20% heat-inactivated FCS, 1 mM L-glutamine, 100 units/mL penicillin, 100 μg/mL streptomycin, 1mM sodium pyruvate and 0.5 mM HEPES (R20). Cells were transferred to wells of 24 well culture plates in a 37°C /5% CO_2_ incubator overnight. The following day, cells were recovered and washed in 10mL R10, supernatants decanted, 1mL of R10 added and cells counted.

RPMI 1640, L-glutamine, HEPES, penicillin/streptomycin, sodium pyruvate and heat-inactivated FCS were all purchased from Sigma-Aldrich (St Louis/Missouri, USA). All laboratories used the same batch of FCS for PBMC isolation and ELISpot assay, which was pretested for acceptable performance in procedures for PBMC isolation, freezing, recovery from frozen and low background (mock) and acceptable PHA/CMV responses in ELISpot. PVDF membrane plates were obtained from Millipore (MSIPS4510; United Kingdom). Anti-human IFN-γ antibody (clone 1-D1K, 1 mg/mL) and biotinylated anti-human IFN-γ antibody (clone 7-B6-1, 1 mg/mL) were purchased from Mabtech, Sweden; peroxidase-avidin biotin complex (ABC) from Vector Laboratories, Burlingame, CA, USA; dimethylformamide (DMF) from VWR International, USA; PHA, 3-amino-9-ethylcarbazole (AEC) tablets (A6926), acetic acid, sodium acetate, 30% hydrogen peroxide (H_2_O_2_) and sterile tissue culture water and phosphate-buffered saline (PBS) were all purchased from Sigma-Aldrich. CEF and CMV pp65 peptides were purchased from Anaspec Inc., CA, USA.

Prior to setting up each ELISpot assay, PVDF plates were treated with 50μL of 70% ethanol for 2 minutes, washed three times with 200 μL/well sterile PBS, coated with 100μL/well of anti-human IFN-γ (1-D1K, 10 μg/mL in PBS) and stored overnight at 4°C. Plates were washed three times with 200 μL/well sterile PBS, blocked for a minimum of 2 hours with 200 μL/well R10 at 37°C, 5% CO_2_ incubator. The blocking medium was removed and 100 μL/well of R10-diluted mock, CEF/CMV peptides (2.25 μg/mL) and PHA (15 μg/mL) were added to their respective wells (following a designated plate plan). 100 μL CMV peptide was added to a designated no cell well. Thawed and overnight-rested PBMC were added at 200,000 cells in 50 μL R10 to each well, with each sample and condition plated in quadruplicate. The plates were incubated for 16-24 hours (37°C, 5% CO_2_). The following day, cells were removed and plates washed six times with 200 μL/well PBS with 0.05% tween (PBS/T) using an automated ELISA washer (Bio-Tek Instruments Inc., Winooski, VT, USA). 100 μL biotinylated anti-human IFN-γ antibody (7-B6-1, 1 μg/mL in PBS with 0.1% BSA) were added for two hours at room temperature (RT). Plates were washed six times as above before addition of 100 μL/well peroxidase avidin-biotin complex (per manufacturer’s instructions), for one-hour at RT. Plates were washed three times with 200 μL/well PBS/T followed by three washes with 200 μL/well PBS. 100 μL/well of 0.45μm filtered AEC substrate (AEC tablet dissolved in 2.5mL DMF, added to 47mL sterile tissue culture water containing 280μL 2M acetic acid and 180μL 2M sodium acetate and finally 25μL H_2_O_2_ added) was added for 4 min at RT. Plates were emptied thoroughly and the reaction stopped under gently-running tap water and the underdrain removed before leaving the plates to dry overnight protected from light. The acceptance criteria for the IFN-γ ELISpot was the mock wells should have less than 10 spots per well and the peptide/media alone (no cells) 5 or fewer spots per well.

#### Data Acquisition and Analysis

Spots were evaluated with an AID reader system (AutoImmun Diagnostika, Germany) with software version 4.0. Each laboratory used the same model of AID reader and defined spot parameters. Responses are expressed as spot-forming cells (SFC) per 10^6^ viable PBMCs as shown in **Figure 6**.

Our main outcomes included 1) the recovery and viability rates of frozen PBMCs, 2) ELISpot results compared to mock, CMV and CEF stimuli and 3) comparison of ELISpot results across laboratories. For each peptide repeated measures, Poisson regression model was fit on background-subtracted count (except mock), with counts from the same volunteer assumed to be correlated. The resulting least squares parameter estimates are presented together with their 95% confidence intervals adjusted for multiple comparisons using the Bonferroni method. Each model included volunteer, laboratory and month. Pair-wise comparisons between laboratories are shown as the ratio of the least squares estimates of mean count with corresponding adjusted (Bonferroni) 95% confidence interval. Statistical significance is defined as a 95% CI for the ratio that excludes unity (i.e., entirely above or below the value 1). Figures 1, 4 and 5 were generated by Graph Pad prism software version 7.01. Other figures and statistical analyses were performed using SAS Version 9.3.

## Results

### 1. Performance in ELISpot assay across 7 laboratories

Each assay included 4 replicates for each peptide. Results based on less than 4 replicates were assumed to be less reliable and excluded from analysis. For each peptide and mock stimulus there were 1751 assays performed of which 50 were excluded (i.e., about 2.9%).

The distribution of mock, CMV and CEF responses across laboratories over time is shown in **Figure 1A & B**. To compare ELISpot data across the 7 laboratories, the response was background-subtracted counts (except mock). The covariates in the model were sample, laboratory and month.

Across the seven laboratories, the geometric mean ELISpot counts (SFC/10^6^ PBMC) were 6 – 10 for Mock, 289 – 438 for CEF and 172 – 266 for CMV (**S1A, S2A & S3A Tables respectively**).

Statistical differences were observed between laboratories in mock counts as shown in **S1B Table** and **Figure 2B** (p=0.0007). For example, the mock count at CLS is estimated to be 1.73 times the mock count at ZEHRP. Also, the mock count at PSF is 0.78 times lower than at UVRI-IAVI. ZEHRP tends to have lower mock counts than all other laboratories.

**Figure 2.**
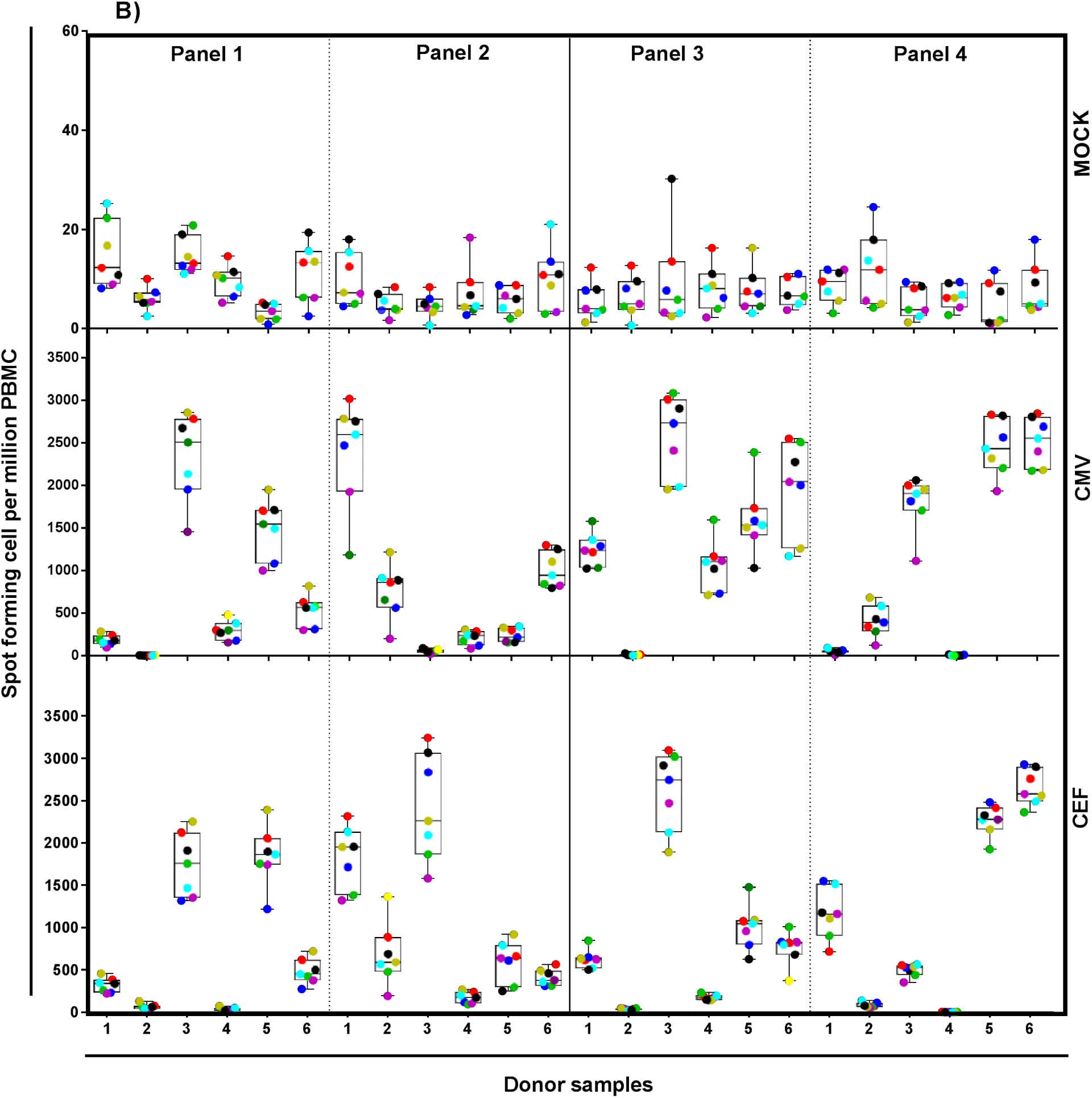
Comparisons of PBMC ELISpot responses: All pair-wise least squares means and their significance, on a natural log scale, for mock, CEF and CMV respectively. For each comparison a line segment, centred at the least squares-means in the pair, is drawn. The segment length corresponds to the projected width of a confidence interval for the least squares mean difference. Segments that fail to cross the 45° reference line correspond to significant least squares mean differences. The graph shows which site pairs are significantly different (blue lines) and which are not (red lines).

When comparing the responses against CEF peptides across laboratories, KEMRI-CGMRC had significantly higher counts than other laboratories (p=0.0045, **S2B Table & Figure 2B).** When data for KEMRI-CGMRC are excluded, the overall difference between laboratories is not statistically significant (p=0.11, **S2C Table**).

When comparing the responses against CMV responses across laboratories, KEMRI-CGMRC again had significantly higher counts than other laboratories (p=0.012, **S3B Table & Figure 2B**). On excluding data for KEMRI-CGMRC the overall difference between laboratories is still statistically significant (p=0.033, **S3C Table**), due to the counts at ZEHRP being lower than CLS and KAVI-ICR.

#### Inter-operator analysis

ELISpot data from 3 operators at KAVI-ICR were compared. Samples from 12 volunteers were analyzed by 3 operators on 2 occasions, each operator analyzing the same set of samples at the same monthly time points. ELISpot counts were obtained for mock and background-subtracted CMV and CEF peptide pools. The covariates in the model were sample, operator and month across the three operators, the geometric mean ELISpot counts (SFC/10^6^ PBMC) were 9 – 12 for Mock, 368 – 393 for CEF and 538 – 598 for CMV (**S4A Table**). The differences between operators were not statistically significant (**S4B Table and Figure 3)**.

**Figure 3.**
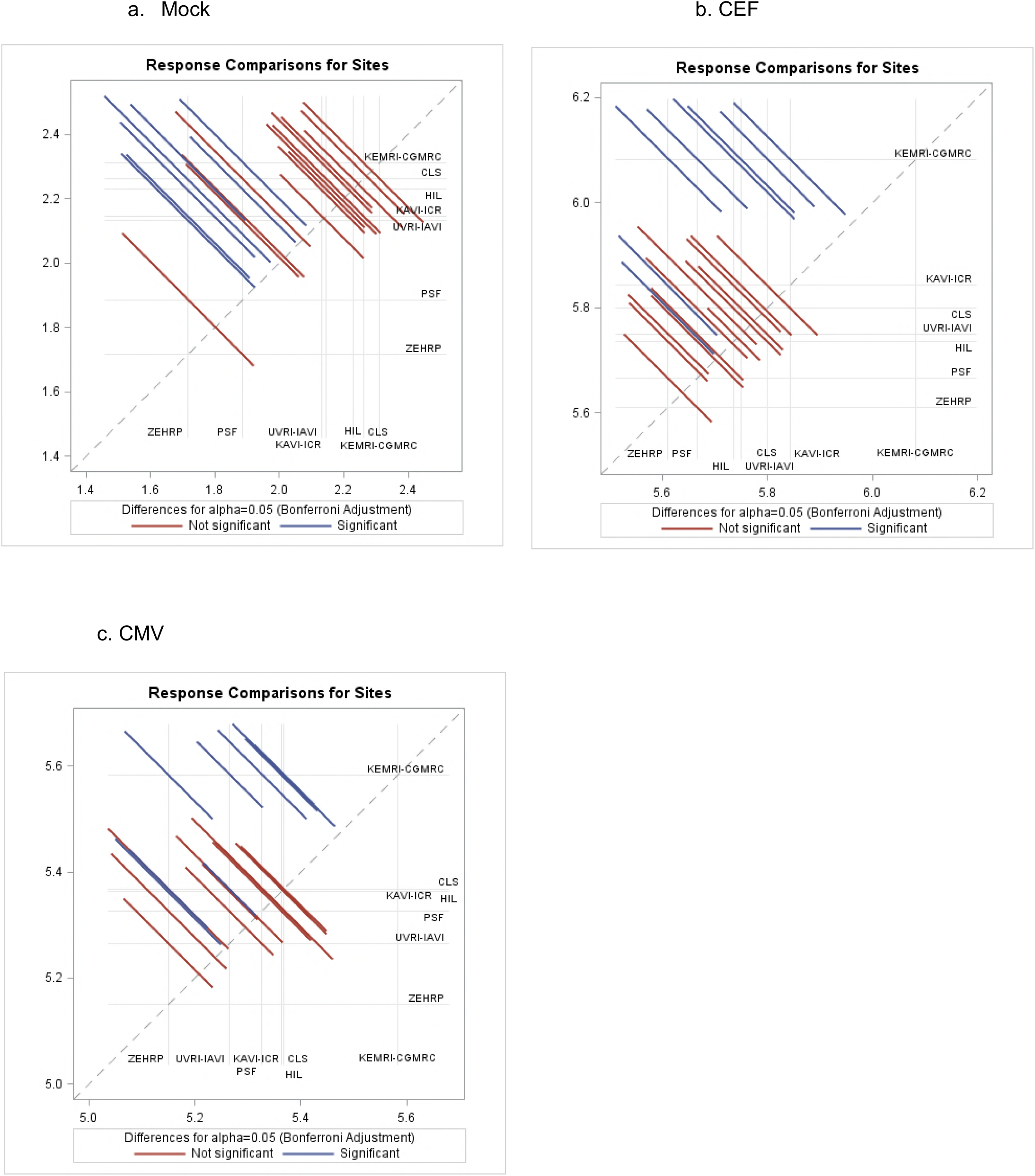
Inter-operator ELISpot comparison from 3 operators at KAVI-ICR: All pair-wise least squares means and their significance, on a natural log scale, for Mock, CEF and CMV respectively. For each comparison a line segment, centred at the lest squares-means in the pair, is drawn. The length of the segment corresponds to the projected width of a confidence interval for the least squares mean difference. Segments that fail to cross the 45° reference line correspond to significant least squares mean differences. None of the pairs of operators are significantly different (all lines cross the 45-degree reference line).

### 2. Viability and cell yield of freshly isolated PBMCs and recovery from frozen PBMCs across 4 laboratories

A total of 1297 PBMCs isolated from clinical trial samples were analysed for cell viability, recovery and cell yield in four laboratories supporting two IAVI-sponsored clinical trials. 1220 of 1297 (94%) freshly-isolated PBMCs had viabilities above 90% with a median of 95% (range 81-100%) while those below 90% had a median of 88% (range 81-90%, **Figure 4A**). Over 96% of these samples had cell yields greater than 0.7×10^6^ PBMC/mL blood, within the pre-defined acceptability criteria with few samples having low cell yield ranging from 0.13-0.56×10^6^ PBMC/mL blood (**Figure 4B)**. A total of 1205 of these samples were tested in ELISpot assay and almost all (99%) had viabilities above 80% following thaw and overnight rest (within acceptability criteria) with only 9 samples having viabilities below 80% ranging from 66 to 78% as shown in **Figure 4C.** Cell recoveries for these samples were above 6.0×10^6^ PBMC/vial (PBMCs were frozen at 10-15×10^6^ PBMC/vial); data were normalized to 10 million cells as shown in **Figure 4D.** For all samples tested, cells were functional in ELISpot assay with over 95% of the samples having mock responses <50 SFC/10^6^ PBMC, PHA>1000 SFC/10^6^ PBMC and a range of CMV responses.

**Figure 4.**
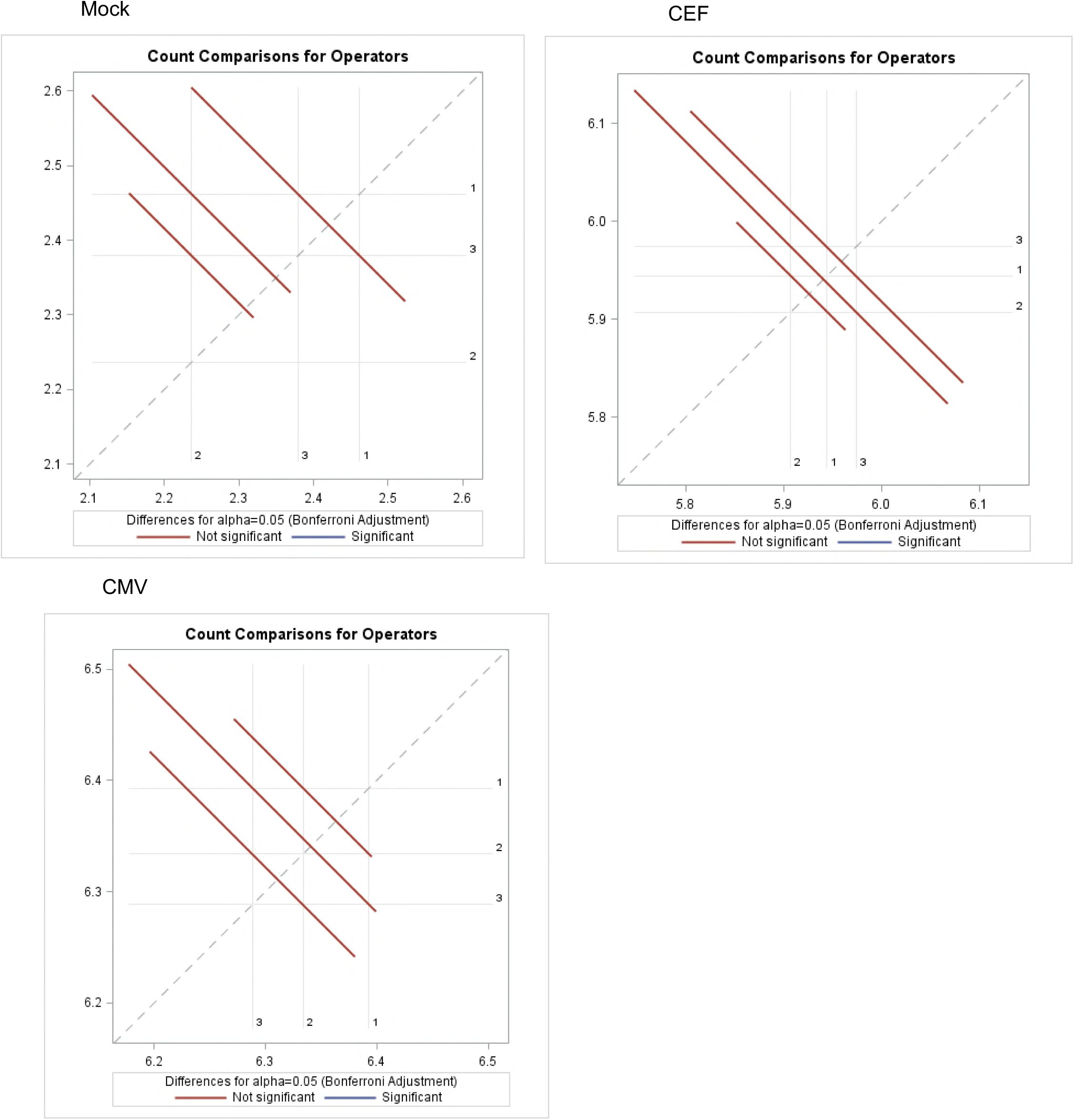
Cell recovery, viability and processing time of clinical trial samples: **A)** viability of freshly isolated PBMC; **B)** cell yield per mL of blood; **C)** viability after overnight rest; **D)** cell recovery of PBMC frozen at 10-15×10^6^ PBMC per vial; data were normalized to 10 million cells; **E)** PBMC processing time. Each point in the scatter plot represents a sample and the lines represent the median with interquartile range. Horizontal lines show the acceptance cut-off.

The length of time from blood draw to sample processing and freezing has been shown to affect the integrity of PBMC (20–22). Nearly all of our samples were processed within 6 hours with 81 (6 %) processed beyond 6 hours (range 6.1-9.5 hours, **Figure 4E & 5E**). To assess the impact of longer processing of these samples, the cell yields and viabilities from fresh blood were analysed together with cell recoveries and viabilities following freezing and thawing. All samples except one had freshly-isolated cell viabilities and cell yields within the acceptable range, that is >90% and >0.7×10^6^ PBMC per mL blood respectively (**Figure 5A & B).** Only one sample had a slightly lower cell yield of 0.57×10^6^ per mL blood (98% viability). Post PBMC freezing cell viabilities ranged from 93-100% and recoveries above 6×10^6^ PBMC/vial in 71/81 (87%) samples (**Figure 5C & D**). We further tested these samples in ELISpot assay to assess their cell functionality and all samples performed well with the mock responses <50 SFC/10^6^ PBMC, PHA responses > 1000 SFC/10^6^ PBMC and a typical range of CMV responses indistinguishable from samples processed within 6 hours, as shown in **Figure 5F.**

**Figure 5.**
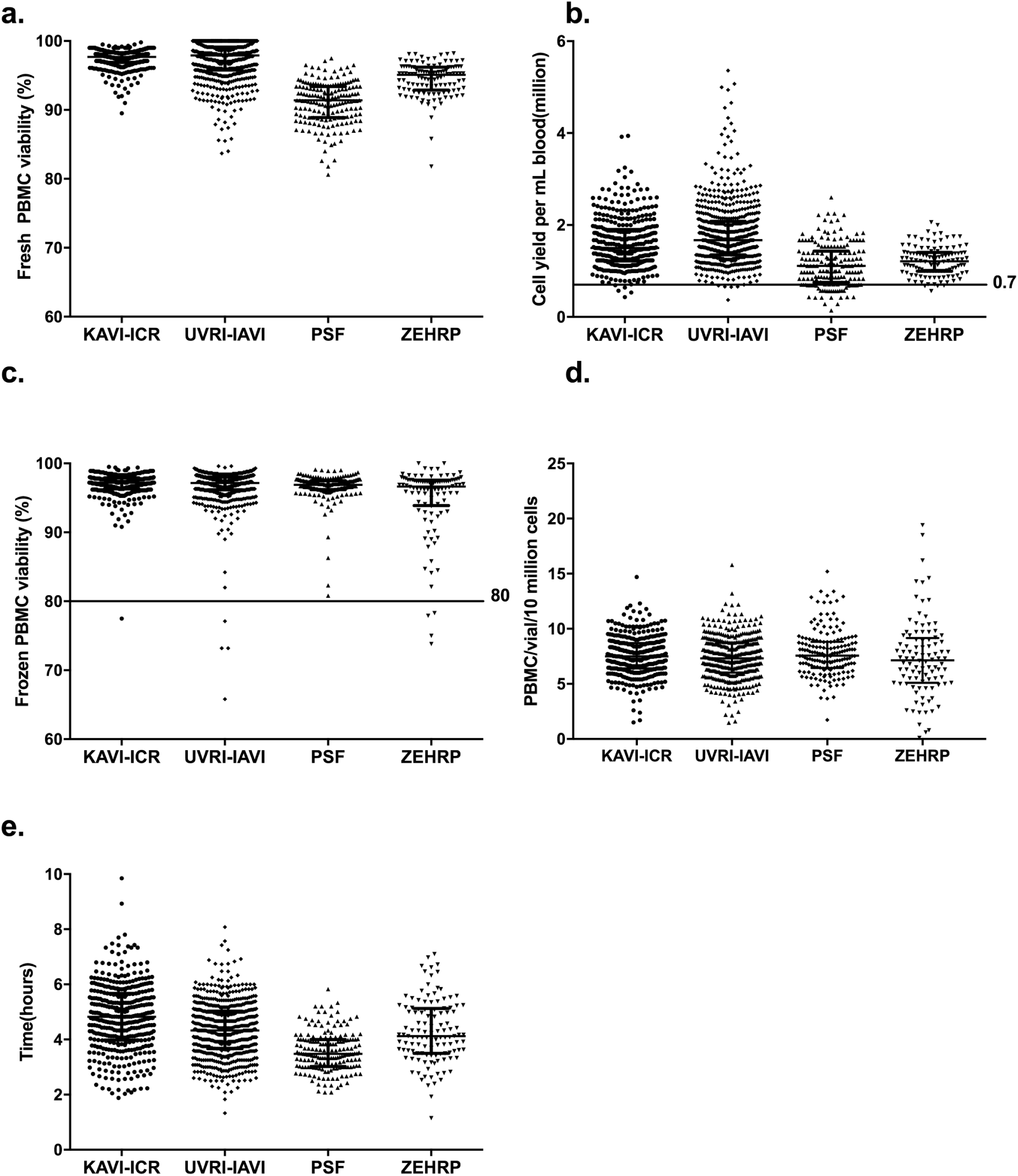
Cell recovery, viability and ELISpot responses of samples processed beyond 6 hours: **A)** viability from freshly isolated PBMC; **B)** cell yield per mL blood; **C)** viability from frozen samples; **D)** cell recovery of PBMC per 10 million cells frozen following thaw and overnight rest; **E)** processing time from blood draw to freezing of PBMC**; F)** ELISpot responses of PBMC tested against mock, PHA and CMV stimuli for PBMCs processed within 6 hours (red) and beyond 6 hours (black). Each point in the scatter plot represents a sample and the median (horizontal line). Horizontal lines represent the acceptance cut-offs.

**Figure 6.**
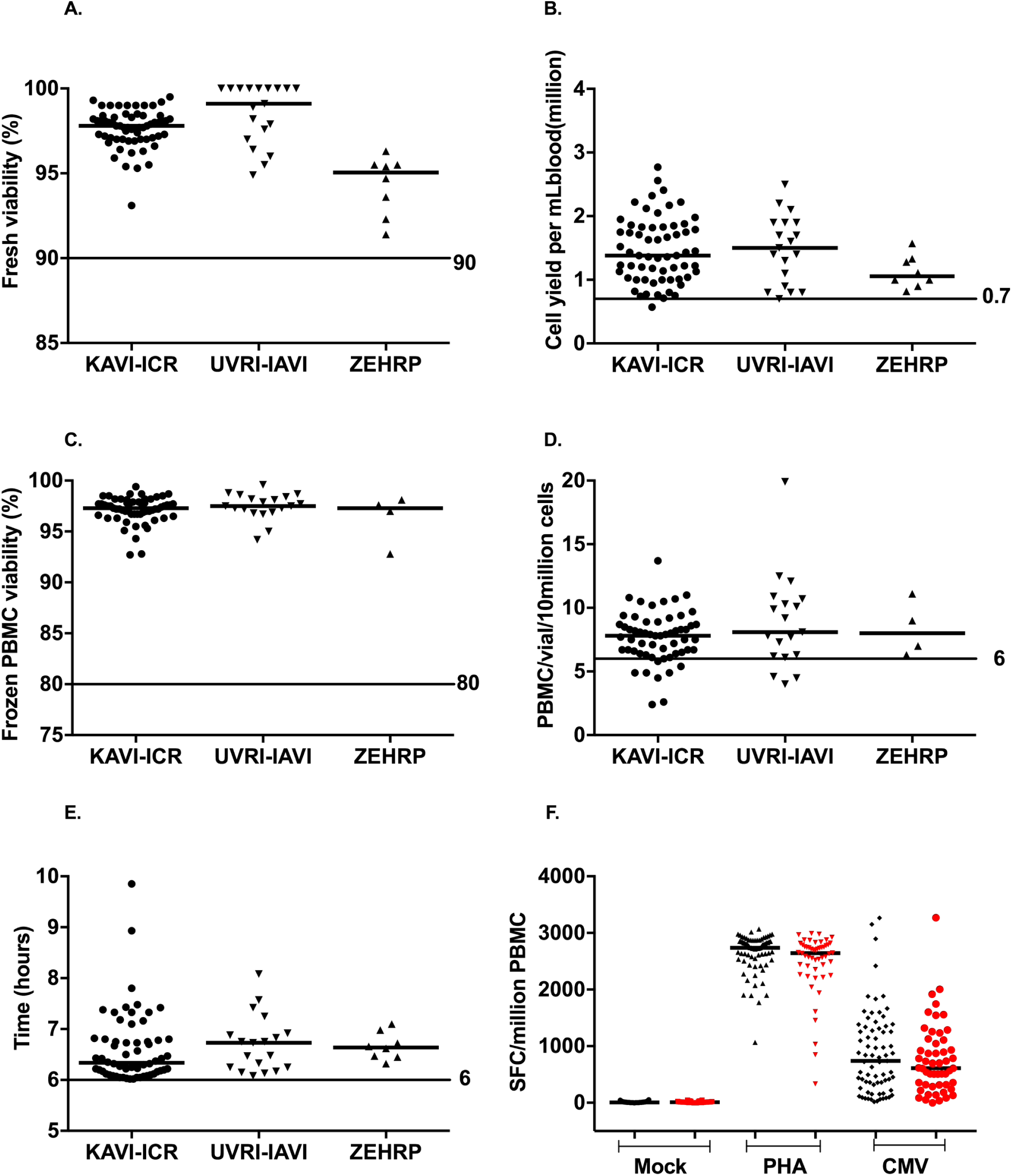
PBMC ELISpot responses against CMV peptides. Representative well images (plate wells C9 to C12) of the CMV responses for sample 3522 performed in quadruplicates across the laboratories. The SFC counted per well are given in each well image.

## Discussion

This report compares data generated across a network of 7 IAVI-supported GCLP-accredited clinical trial laboratories based in Africa and Europe. These laboratories were assessed on ELISpot proficiency testing and PBMC processing.

ELISpot proficiency data were analysed and compared across and within sites. Acceptability criteria for mock and PHA controls and CMV and CEF responses had to be clearly positive or negative during pre-testing. We found that all laboratories correctly detected responses against CMV and CEF peptide stimuli with the exception of a few sporadic higher data points in mock stimulus which was seen across laboratories. As expected even for technically competent laboratories there were occasional discrepant data points.

In this report we have demonstrated that ELISpot data for CEF and CMV responses from 5 laboratories were not significantly different and were overall comparable. Four of these laboratories were actively performing ELISpot analysis in support of IAVI clinical trials whilst a fifth laboratory (CLS) performed ongoing ELISpot analyses as part of establishing the proficiency scheme and in training of staff at other laboratories. We did observe significantly different data for CEF and CMV responses from 2 laboratories that were not routinely conducting ELISpot assay in support of IAVI clinical trials when compared to the other laboratories. Staff at these laboratories only performed ELISpot analyses as part of the proficiency program described in this report and therefore would have less ongoing technical experience in ELISpot analysis compared to staff at laboratories with active participation in clinical trials. To mitigate this, staff at the 2 laboratories were retrained and competency assessed. This highlights the need for review of staff retraining and continual monitoring of laboratories’ performance with trouble-shooting and staff training and re-training as required, especially for laboratories taking on new activities or trials or where laboratories have not performed certain techniques in a trial setting for some time. However, although the ELISpot responses observed at these 2 laboratories were statistically significantly different, the range of estimated least squares mean counts across the 7 laboratories was not high with 274 to 438 for CEF (**S2A Table**) and 172 to 266 SFC/10^6^ PBMC for CMV (**S3A Table**). Statistically significant differences in mock values between laboratories were apparent which may be expected as in effect the vast majority of mock responses were close to zero. Across the seven laboratories, the geometric mean ELISpot mock counts were 6 – 10 SFC/10^6^ PBMC (**S1A Table**). High variability of low T cell responses has been reported previously (23).

Operator-dependent variability in ELISpot is a known phenomenon (24) and we assessed this in this report. It was not possible to analyse inter-operator variability at all laboratories as some laboratories had either a lone operator throughout, or a change of operators during the study period. However, we report on one laboratory with 3 operators performing the ELISpot assay on a rotational basis. All operators detected correctly the expected responses for CMV and CEF stimuli. Their data were highly correlated and variability in data points was not significantly different.

Achieving accurate and reliable results when assessing the immunogenicity of vaccine candidates, especially for multi-site clinical trials, is essential. In order to achieve this, samples must be processed according to standardised SOPs following GCLP-guidelines for data integrity. PBMC processing in four clinical trial laboratories were analysed for processing time from blood draw to start of freezing, cell yield and viability and post-freezing viability and recovery. We report that the vast majority of freshly isolated PBMCs had viabilities and cell yields within the acceptable range across all laboratories.

Proper freezing and storage of samples is critical in preserving cell integrity and functionality (25). In this report we assessed the integrity of PBMCs processed and frozen at the laboratories. Cells were thawed at the HIL (Central repository lab) for ELISpot testing and nearly all samples had cell viabilities and recoveries within the acceptance criteria, with cell functionality demonstrated by good performance in ELISpot assay, therefore demonstrating the competency of laboratories in isolation, freezing, storing and shipping of PBMC samples.

PBMCs processed beyond 8 hours have been shown to have reduced cell viability and compromised cell functionality (26). Here, we report that the majority of samples were processed within 6 hours with the exception of few samples processed beyond 6 hours, mainly due to delayed delivery to the laboratories from some clinics located up to 50 miles away. For samples processed beyond 6 hours, corrective and preventive action (CAPA) reports were written to minimise or prevent recurrence where possible and monitored on a monthly basis. Cell yields, viabilities and recoveries of these samples were assessed to determine the impact of longer processing on their integrity. All samples performed well in ELISpot with responses to Mock, CMV, CEF and PHA stimuli being in the expected ranges with data similar to samples processed within 6 hours. Therefore, we show that PBMCs processed longer than 6 hours (up to 9 hours) are still viable and functional in ELISpot assay and similar to what other groups have shown (20–21).

Participating laboratories are audited regularly for GCLP compliance by internal and external independent auditors. The audit covers SOPs, ELISpot and flow cytometry proficiency, external quality assurance programs and data integrity. The GCLP audit by an external auditor from Qualogy Ltd, UK is conducted annually with an accreditation certificate issued to compliant laboratories.

In conclusion, we have demonstrated that using standardised SOPs, equipment and reagents and working in a GCLP compliant laboratory, clinical trial laboratories located in Africa and Europe can process clinical trial samples and maintain cell integrity and functionality through ELISpot testing, producing comparable and reliable data.

## Acknowledgments

We wish to acknowledge the support from the University of California, San Francisco’s (UCSF) International Traineeships in AIDS Prevention Studies (ITAPS), U.S. NIMH, R25 MH064712 under which this manuscript was written. We thank UCSF faculty staff for helpful discussions, Matt Price and Kathy Crisafi of IAVI for manuscript review and laboratory technicians for experimental assistance. We also thank all IAVI-Clinical Research Centres’ Principal Investigators for overseeing the laboratories. This work was funded in part by IAVI and made possible by the support of the United States Agency for International Development (USAID) and other donors. The full list of IAVI donors is available at www.iavi.org. The contents of this manuscript are the responsibility of the authors and do not necessarily reflect the views of USAID or the US Government.

